# riboCIRC: a comprehensive database of translatable circRNAs

**DOI:** 10.1101/2021.03.03.433665

**Authors:** Huihui Li, Mingzhe Xie, Yan Wang, Ludong Yang, Zhi Xie, Hongwei Wang

**Affiliations:** State Key Laboratory of Ophthalmology, Zhongshan Ophthalmic Center, Sun Yat-sen University, Guangzhou, China

**Keywords:** CircRNAs, Translatable circRNAs, Ribosome profiling, circRNA-encoded peptides

## Abstract

riboCIRC is a translatome data-oriented circRNA database specifically designed for hosting, exploring, analyzing, and visualizing translatable circRNAs from multi-species. The database provides a comprehensive repository of computationally predicted ribosome-associated circRNAs, a manually curated collection of experimentally verified translated circRNAs, an evaluation of cross-species conservation of translatable circRNAs, a systematic *de novo* annotation of putative circRNA-encoded peptides, including sequence, structure, and function, and a genome browser to visualize the context-specific occupant footprints of circRNAs. It represents a valuable resource for the circRNA research community and is publicly available at http://www.ribocirc.com.

## Background

Circular RNAs (circRNAs) are an abundant class of covalently closed endogenous RNA molecules generated by back-splicing of pre-mRNAs. Recent advances in computational analysis and high-throughput RNA sequencing (RNA-seq) have unveiled a detailed view of circRNA biogenesis, regulatory mechanisms and cellular functions [1]. With the development of various computational and experimental approaches to effective identification of circRNAs, many dedicated databases for circRNAs were constructed, such as circBase and circAtlas for vertebrate circRNAs [2,3], CSCD and TSCD for disease/tissue-specific circRNAs [4,5], and Circ2Disease and Circ2Traits for circRNA-disease associations [6,7]. These transcriptome data-oriented databases provide essential information about circRNAs, facilitating the current understanding of circRNAs related to their biological importance and clinical relevance. It becomes increasingly clear that circRNAs can regulate multiple biological processes via a variety of mechanisms. For instance, circRNAs can act as ‘sponges’ or ‘decoys’ for microRNAs or RNA-binding proteins to modulate gene expression or mRNA translation [8–10].

circRNAs are generally considered as ‘non-coding’ elements; however, circRNAs can in fact serve as templates for protein translation. Using ribosome profiling (Ribo-seq) that enables genome-wide investigation of *in vivo* translation at a sub-codon resolution [11,12], a subset of circRNAs have recently been identified to be associated with translating ribosomes [13,14]. Furthermore, by performing *in vivo* and *in vitro* translation assays, circRNAs have been shown to enable cap-independent translation and generate functional proteins. Of these proteins, some have been demonstrated to play vital roles under a number of pathophysiological conditions, such as muscle-enriched circRNA circ-ZNF609 [10] and brain-ubiquitously expressed circRNA circAβ-a [15]. In addition, several mechanisms for circRNAs translation have been proposed. For instance, internal ribosome entry site (IRES)- and N6-methyladenosines (m6A)-mediated cap-independent translation initiation are potential mechanisms for circRNA translation [16,17].

Although translation of circRNAs has attracted considerable attention and a large number of the Ribo-seq datasets have been generated in the past several years [18], there is no translatome data-oriented database that aims to provide direct *in vivo* translation evidence for multi-species circRNAs to date. To fill the gap, we analyzed the 3,168 publicly available Ribo-seq and 1,970 matched RNA-seq datasets from 314 studies covering 21 various species to determine the prevalence of circRNA translation. We further provided a dedicated multi-species translatable circRNA database, riboCIRC, towards a comprehensive repository of computationally predicted and experimentally verified translatable circRNAs. Overall, the riboCIRC database provides an important resource for the circRNA research community and can serve as a useful starting point for further investigation of the details of circRNA function and their involvement in cellular processes and diseases.

## Construction and content

The backend of riboCIRC is powered by MySQL and accessed using the PHP framework as the middleware. The MySQL includes three large tables: the first table stores all information about the available computationally predicted ribosome-associated circRNAs in the database; the second table stores all information about the manually curated experimentally verified circRNAs; and the third table stores all annotation information about the putative circRNA-encoded peptides. The front-end of riboCIRC is a multi-page web application built using HTML, JavaScript, and CSS code that consists of numerous pages with static information, such as text and image. The entire database is deployed and running on Amazon AWS EC2 platform. The current version of riboCIRC records a total of 2,247 computationally predicted ribosome-associated circRNAs and 216 experimentally verified translated circRNAs from six different species. The web application of riboCIRC is designed for hosting, exploring, analyzing and visualizing these translatable circRNAs (**Figure 1**). The home page provides general information about riboCIRC and quick links to access translatable circRNAs, circRNA-encoded peptides, and footprint visualization of circRNAs. Other primary features of riboCIRC can be accessed using various buttons of the website.

**Figure 1.**
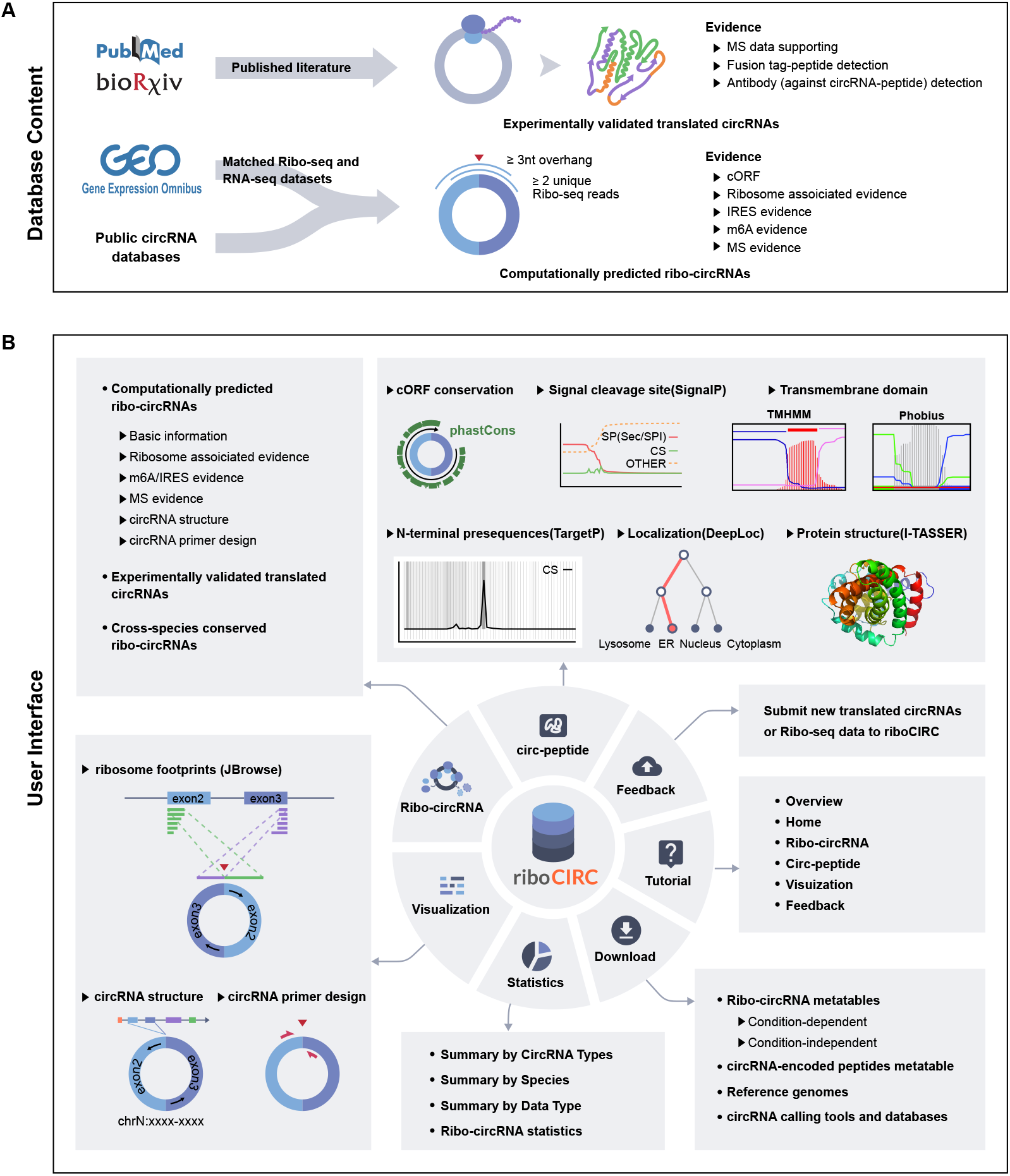
Overview of the riboCIRC database. (a) Schematic view of the database content. (b) Schematic view of the database interface.

### Data collection and preprocessing

We collected 3,168 publicly available Ribo-seq datasets and 1,970 matched RNA-seq datasets of the same samples from 314 studies covering 21 species, including *Arabidopsis, Caenorhabditis elegans, Caulobacter crescentus, Cryptococcus neoformans*, Chinese hamster, *Drosophila, Escherichia coli, Halobacterium salinarum*, human, mouse, *Plasmodium falciparum, Pseudomonas aeruginosa*, rat, *Saccharomyces cerevisiae, Salmonella enterica, Schizosaccharomyces pombe, Streptomyces coelicolor, Staphylococcus aureus, Trypanosoma brucei, Vibrio vulnificus*, and zebrafish (see **Additional file 1: Table S1**). After downloading the raw data files from the NCBI SRA database [19], we applied a unified pipeline to perform preprocessing of the Ribo-seq and RNA-seq data. Briefly, the 3’-end adapters were clipped using Cutadapt (version 1.8.1) [20]; Low-quality bases were trimmed using Sickle (version 1.33) [21]; and the retained reads that mapped to rRNAs or tRNAs were removed.

### Detection of transcribed and ribosome-associated circRNAs

We combined three different detection tools to identify transcribed circRNAs in each RNA-seq dataset, namely, CIRCexplorer2, CIRI2, and DCC [22–24]. The full-length sequence of each identified circRNA was assembled by the CIRI-full pipeline [25] or extracted from the circAtlas database [2,3] when RNA-seq data were unavailable. Taking advantage of these full sequences, we generated a pseudo circRNA reference for each species by initial extraction of the 23-base pair (bp) sequences on either side of the backsplice junction (BSJ) site of each transcribed circRNA with subsequent concatenation of the two-sided sequences. To identify ribosome-associated circRNAs (ribo-circRNAs), we first eliminated sequence reads corresponding to nonribosomal RNA-protein complexes in each Ribo-seq dataset using Rfoot (version 1.0) [26], considering that ribosomes are not specifically selected during the biochemical isolation procedure of ribosome profiling experiment. After removal of footprints from nonribosomal complexes, all the ribosome-protected footprints were then mapped with Tophat2 (version 2.1.1) [27] to the corresponding linear reference genome, and further the resulting unmapped.bam files were remapped to the pseudo circRNA reference using Tophat2 (version 2.1.1) [27] with default parameters except N, which was set to 0 (the default is 2). Finally, a circRNA was defined to be associated with translating ribosomes only when it met all of the following three criteria simultaneously: (1) at least two unique backsplice junction-spanning Ribo-seq reads, (2) a minimum read-junction overlap of three nucleotides (nt) on either side of the backsplice junction site, and (3) a typical range of read lengths of 25-35 nt (see **Additional file 2: Figure S1**).

Two different strategies were here used to characterize ribo-circRNAs: (1) condition-dependent detection for Ribo-seq and perfectly matched RNA-seq datasets and (2) condition-independent detection for previously reported circRNAs and Ribo-seq datasets. The former strategy was applied to the initial genome-wide characterization of transcribed circRNAs using 1,922 RNA-seq datasets with subsequent examination of ribosome associations of these circRNAs using 1,970 Ribo-seq datasets from the same samples. In total, 278 out of the 91,143 transcribed circRNAs were identified as ribo-circRNAs, involving four different species (*Drosophila*, human, mouse, and rat). The latter strategy was applied to the systemic examination of ribosome associations of the circRNAs reported in the public databases using 3,168 Ribo-seq datasets. To accomplish this task, we selected nine out of the 18 examined public circRNA databases, including circAtlas, circBank, circBase, CIRCpedia, circRNADb, CSCD, exoRBase, TSCD and Circ2Disease [2–6,28–31], and obtained 1,411,865 unique circRNAs after conversion of their coordinates using LiftOver [32]. Notably, the other public circRNA databases were excluded from this analysis due to lack of a batch download link, incomplete annotation or inaccessible webpage (see **Additional file 3: Table S2**). Among these well-documented circRNAs, a total of 1,969 circRNAs were finally identified as ribo-circRNAs, involving six different species (*C. elegans, Drosophila*, human, mouse, rat, and zebrafish).

### Cross-species conservation analysis of translatable circRNAs

To evaluate translatable circRNA conservation among different species, we first annotated the parental genes of ribo-circRNAs using the GTF files, and then identified orthologous gene pairs expressing these circRNAs using a pairwise orthologous gene list downloaded from the OMA orthology database (http://omabrowser.org) [33]. After that, we extracted 50-bp fragments on either side of the ribo-circRNA BSJ site from the reference genome and further concatenated both fragments to represent the ribo-circRNA BSJ sequence. Next, all ribo-circRNA BSJ sequences in one species were aligned to those of the other species using BLAT with default parameters [34], followed by a reciprocal best hit strategy to find the orthologous ribo-circRNAs. Finally, a pair of conserved ribo-circRNAs were defined based on their sequence alignment length ≥ 80 and alignment bit-score ≥ 150.

### Prediction of circRNA-derived ORFs

We predicted putative circRNA-derived ORFs (cORFs) for each ribo-circRNA using the cORF_prediction_pipeline with some modifications [13]. Briefly, the full-length sequence of each ribo-circRNA was retrieved and multiplied four times to allow for rolling circle translation. All cORFs beginning with an AUG initiation codon were identified separately for each circRNA, and further filtered based on the requirements of a minimum length of 20 amino acids (aa) and of spanning the backsplice junction site. Notably, those cORFs terminating without an in-frame stop codon were defined as INF (infinite)-cORFs, representing that the corresponding circRNAs could be translated via a rolling circle amplification mechanism. Finally, only the longest cORF was kept for each one of the three reading frames, considering that circRNA with a long ORF would have a better chance of undergoing translation.

### Annotation of IRES elements and m6A sites in circRNAs

Given that previous studies have shown the ability of IRES elements and m6A modification to drive circRNA translation [13,16], we predicted potential IRES elements and m6A sites in circRNAs by using publicly available IRES sequences and m6A modification data. To identify IRES elements in circRNAs, we extracted experimentally validated IRES sequences from the IRESbase database [35], and then aligned them to circRNA sequences using BLASTN (version 2.7.1+) [36] with at least 80% sequence identity and a cutoff 30 nucleotides alignment length. To identify potential m6A sites in circRNAs, we extracted m6A modification peaks detected by three different peak calling tools (exomePeak, MeTPeak, and MACS2) from the REPIC database [37], followed by aligning them to circRNA sequences and the presence of m6A consensus motif ‘RAC’ (where R is any purine) in the aligned positions.

### Annotation of cORF-encoded peptides

We constructed a semi-automated bioinformatic workflow system to perform *de novo* annotation of all putative cORF-encoded peptides, including sequence conservation, transmembrane topology, signal cleavage site, subcellular localization, folding structure, potential function, etc. Specifically, sequence conservation of each putative cORF-encoded peptides was computed by an in-house Python script based on the phastCons score files at the University of California Santa Cruz (UCSC) [38]. The presence or absence of the signal peptide cleavage sites was predicted by SignalP (version 5.0b) with default parameters [39]. Transmembrane helical topology was predicted by TMHMM (version 1.1) [40] and Phobius (version 1.01) [41] with default parameters. The N-terminal presequences, such as signal peptide (SP), mitochondrial transit peptide (mTP), chloroplast transit peptide (cTP) or thylakoid luminal transit peptide (luTP), were predicted by TargetP (version 2.0) [42] with default parameters. Subcellular localization was predicted by DeepLoc (version 1.0) [43] with default parameters, which can differentiate between 10 different localizations, including nucleus, cytoplasm, extracellular, mitochondrial, cell membrane, endoplasmic reticulum, chloroplast, Golgi apparatus, lysosome/Vacuole, and peroxisome. The 3D structure was predicted by I-TASSER (version 5.1) [44] that also provided other information, such as secondary structure, solvent accessibility, normalized B-factor and Top 10 threading templates.

### Detection of cORF-encoded peptides by mass spectrometry

We used public proteomics data to find protein evidence of putative cORF-encoded peptides. Briefly, the raw files were of 26 datasets downloaded from the PRIDE database [45] (see **Additional file 4: Table S3**) and analyzed using MaxQuant software (version 1.6.15.0) [46] against a custom-tailored database separately for each species (the respective size for human: n=22,113; mouse: n=18,308; rat: n=8,159; *Drosophila:* n=3,629; *C.elegans:* n=4,156; and zebrafish: n=3,165), which combined all documented sequences from UniProt/Swiss-Prot with additional sequences derived from circRNA translation, based on the target-decoy strategy (Reverse) with the standard search parameters with the following exceptions: (1) the peptide-level FDR was set to 5%, and the protein-level FDR was excluded; (2) the minimal peptide length was set to seven amino acids; and (3) a maximum of two missed cleavages were allowed. For each search, fixed modifications and variable modifications were customized according to different proteomics data. In total, 719 cORF-encoded proteins from 669 circRNAs were evidenced by at least one unique junction-spanning peptide.

### Primer design and structure representation of circRNAs

Based on the sequence of each circRNA, we performed circRNA-specific primer design. Divergent primer sets spanning the backsplice junction sequence were generated using circtools [47]. The graphical representations of circRNAs and their linear host transcripts were constructed using circView [48].

### Collection of experimentally verified translated circRNAs

Experimentally verified translated circRNAs were manually curated from the literature. To accomplish this task, we searched the PubMed literature database using the keyword ‘(circRNA [MeSH terms]) AND (translation [MeSH terms])’ and the bioRxiv preprint server using the keyword ‘(“circRNA”+”translation”)’, and found a total of 65 relevant published or preprint references. After retrieving the full text of these references, we reviewed the studies to manually collect the circRNA entries, which generated the peptides and were validated by various experiments. Strict screening identified 216 translated circRNAs with mass spectrometry-derived detection of the corresponding peptides, tag-peptide fusion system detection, or/and antibody (against circRNA-peptide) detection evidence and incorporated all information into the riboCIRC database. Additional basic information on these experimentally verified circRNAs was also collected, including the circRNA name, circBase id, genomic coordinates, strand, host gene, transcript, species/condition, circRNA-encoded peptide sequence, peptide length, experimental method, and reference information.

## Utility and discussion

### Exploration of translatable circRNAs

The ribo-circRNA page provides a comprehensive repository of translatable circRNAs, including computationally predicted ribosome-associated circRNAs and experimentally verified translated circRNAs. Users can click the ‘Ribo-circRNA’ button on the navigation bar and then select one of the dropdown-menu options (including ‘Computationally predicted ribo-circRNAs’, ‘Experimentally verified translated circRNAs’, and ‘Cross-species conserved ribo-circRNAs’) for a quick query.

Selection of ‘Computationally predicted ribo-circRNAs’ returns the result page containing all predicted ribo-circRNAs identified using Ribo-seq data, including 1,969 condition-independent and 278 condition-dependent ribo-circRNAs. Brief descriptions of these circRNAs are shown in this results page, including riboCIRC id, chromosome position, best transcript, host gene symbol, and circRNA length. A built-in search box can narrow the results down to a particular subject by entering additional search terms. Furthermore, clicking the riboCIRC id in the second column opens a separate page for every circRNA that displays detailed information on the matching circRNA, including cORF annotation such as the location of the junction-spanning cORF in the genome, total number of junction-spanning footprints, unique number of junction-spanning footprints, translation conditions, involved dataset, cORF sequence, cORF-encode peptide, and length of cORF-encode peptide, evidence for translation of circRNAs such as IRES element, m6A site, and mass spectrometric proof, graphical representation of the linear and circular RNA structure, and designed circRNA primer sets. Clicking the chromosome position in the third column opens a separate page for visualization the host gene track, genomic features and aligned junction-spanning ribosome footprints of the circRNA in the JBrowse [49]. Selection of ‘Experimentally verified translated circRNAs’ returns the result page with all 216 experimentally verified translated circRNAs that have been validated by various experiments to generate peptides. All information collected on these circRNAs is shown with some additional relevant information on the validated circRNAs accessible via the hyperlinks provided on the result page. In addition, selection of ‘Cross-species conserved ribo-circRNAs’ returns the result page with all cross-species inference of conserved translatable circRNA pairs that involve a total of 90 evolutionarily conserved ribo-circRNAs.

### Comprehensive analysis of circRNA-encoded peptides

The circ-peptide page provides a systematic annotation of putative circRNA-encoded peptides, including their sequence, structure and function. Users can click the ‘Circ-peptide’ button on the navigation bar to quickly browse the putative circRNA-encoded peptides. A dropdown menu shows a list of the available circRNA-encoded peptide options, and users can select one of the options to retrieve additional information, including basic information on the given peptide (sequence, and conservation), summary of peptide characteristics (signal cleavage site, transmembrane domain, and N-terminal presequence), and location and topology of the peptide (subcellular localization, secondary structure, and structural conformation).

### Intuitive visualization of ribosome-associated circRNAs

The visualization page provides an intuitive view of ribo-circRNAs, including visualization of the host gene track, genomic features and aligned junction-spanning ribosome footprints of the circRNAs. Users can click the ‘Visualization’ button on the navigation bar to visualize the data on the features of ribosome-associated circRNAs in JBrowse browser [49] embedded in the result page. A cascading dropdown menu consists of three independent selection dropdown buttons for quick navigation to a circRNA of interest, interactive exploration of the data, and intuitive comparison of the data originating from various datasets.

### Data download, statistics, user guide, and feedback

The download page provides access to a convenient tabular data format. Users can click the ‘Download’ button on the navigation bar to easily access the data. Tabular list of metadata for all computationally predicted ribo-circRNAs, circRNA sequences, nucleotide sequences of all cORFs and their corresponding protein sequences, as well as customized protein sequence databases for proteomics search can be freely downloaded for nonprofit and academic purposes. In addition, the Ribo-circRNA page also provides the download buttons for download of computationally predicted ribosome-associated or experimentally verified translated circRNAs in various formats, including JSON, XML, CSV, TXT, SQL and MS-Excel. The statistics page provides a summary that summarizes the data in all accessible records of the database. The tutorial page provides step-by-step instructions for users to familiarize themselves with the database. The feedback page provides a feedback form for translatable circRNAs and Ribo-seq datasets, making it easy for users to provide feedback.

### Potential Limitation

It should be noted here that traditional approaches based on the properties of active translation such as three-nucleotide periodic subcodon pattern are not feasible for identifying active translated circRNAs due to the difficulty in distinguishing circular and linear Ribo-seq reads. In this database, we also adopt a similar strategy of translatable circRNA detection as previously described [13,14], where ribosome-associated circRNAs were identified only by Ribo-seq reads spanning a head-to-tail splice junction. However, computational prediction through this strategy does not necessarily mean that the circRNA is being actively translated into a detectable micropeptide, even though it is associated with translating ribosomes. This database is just a starting point for bench scientists and computational biologists to pursue translatable circRNAs. The translated details of individual circRNAs still have to be further experimentally validated.

### Future directions

In the future, riboCIRC will be periodically updated. Increasing availability of new high-throughput Ribo-seq data will be used to characterize the putative translation of circRNAs and expand the size of computationally predicted ribo-circRNAs. We will continue to fill the database with new reported experimentally verified translated circRNAs. In addition, we will continue to extend our collection of public proteomics data and to further enhance the identification rate of cORF-encoded peptides. These additions are anticipated to enhance efficiency of the applications of riboCIRC in the circRNA research community.

## Conclusions

To the best of our knowledge, riboCIRC is the first database for hosting, exploring, analyzing, and visualizing translatable circRNAs for multi-species. The database provides a comprehensive repository of computationally predicted ribo-circRNAs, together with multiple lines of evidence supporting their translation, and experimentally verified translated circRNAs. It also provides an evaluation of cross-species conversed translatable circRNAs, a systematic functional annotation of the putative circRNA-encoded peptides, a flexible visualization framework for ribosome-associated circRNAs, and a user-friendly web interface for easy data access and exploration. Thus, riboCIRC will serve as a valuable resource for bench scientists and computational biologists to explore translatable circRNAs and to drive functional investigation of the circRNA translation.

## Ethical approval and consent to participate

Not applicable.

## Consent for publication

Not applicable.

## Availability of data and materials

riboCIRC is available at http://www.ribocirc.com to all users without any login or registration restrictions. All public Ribo-seq, RNA-seq, and mass spectrometry datasets used during the current study are available in **Additional file 1: Table S1 and Additional file 4: Table S3**. All translatable circRNAs can be downloaded from the riboCIRC data download page.

## Authors’ contributions

H.W.W. and Z.X. directed the project and wrote the manuscript. H.H.L, Y.W. and L.D.Y. performed the data analyses and result presentation. H.H.L. and M.Z.X. designed and constructed the database. All named authors read and approved the final manuscript.

## Acknowledgments

We would like to thank our team members Congying Chen and Liang Yi for their assistance in collecting translated circRNAs. We also thank the support from the Center for Precision Medicine at Sun Yat-sen University.

## Additional files

**Additional file 1: Table S1.**
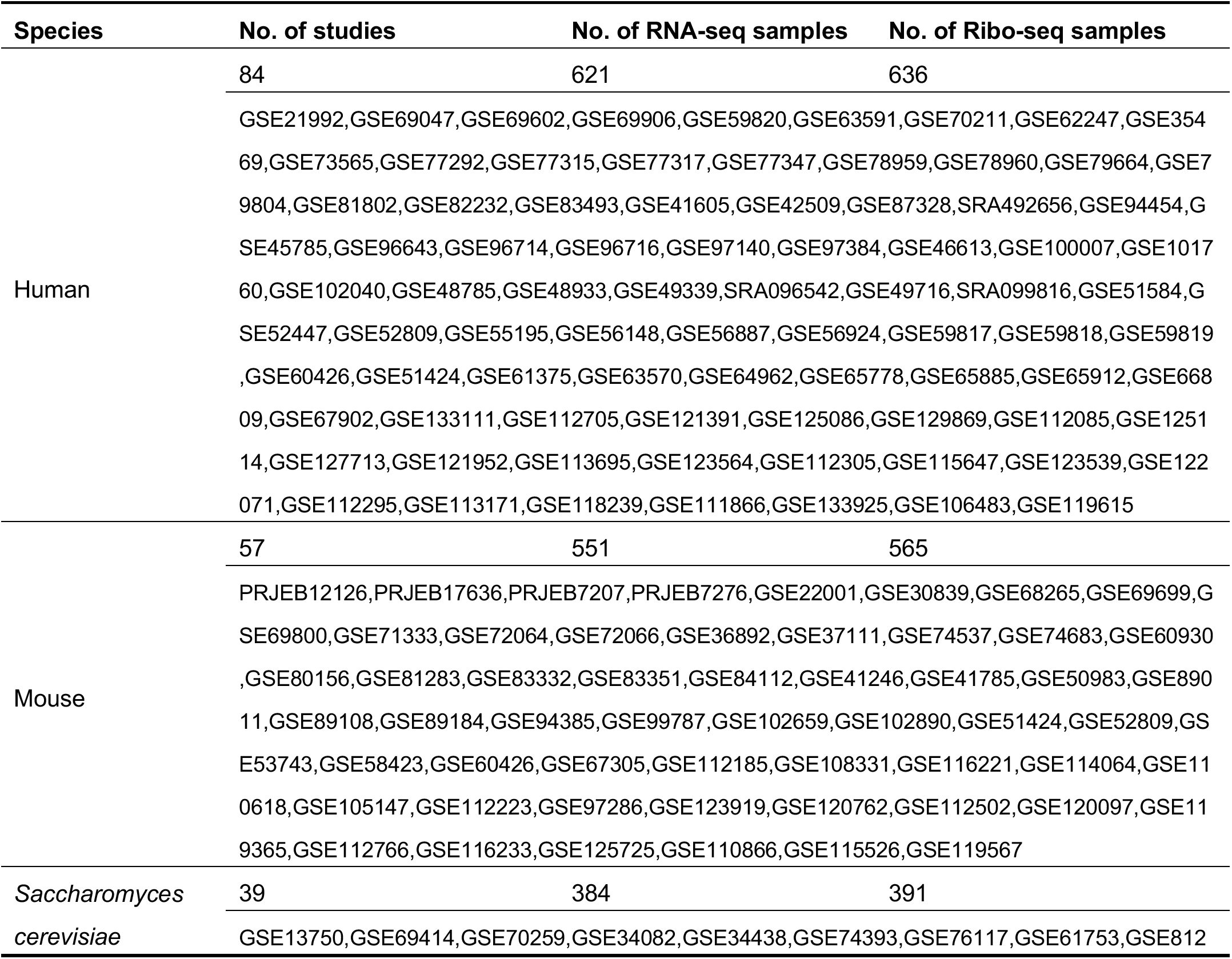
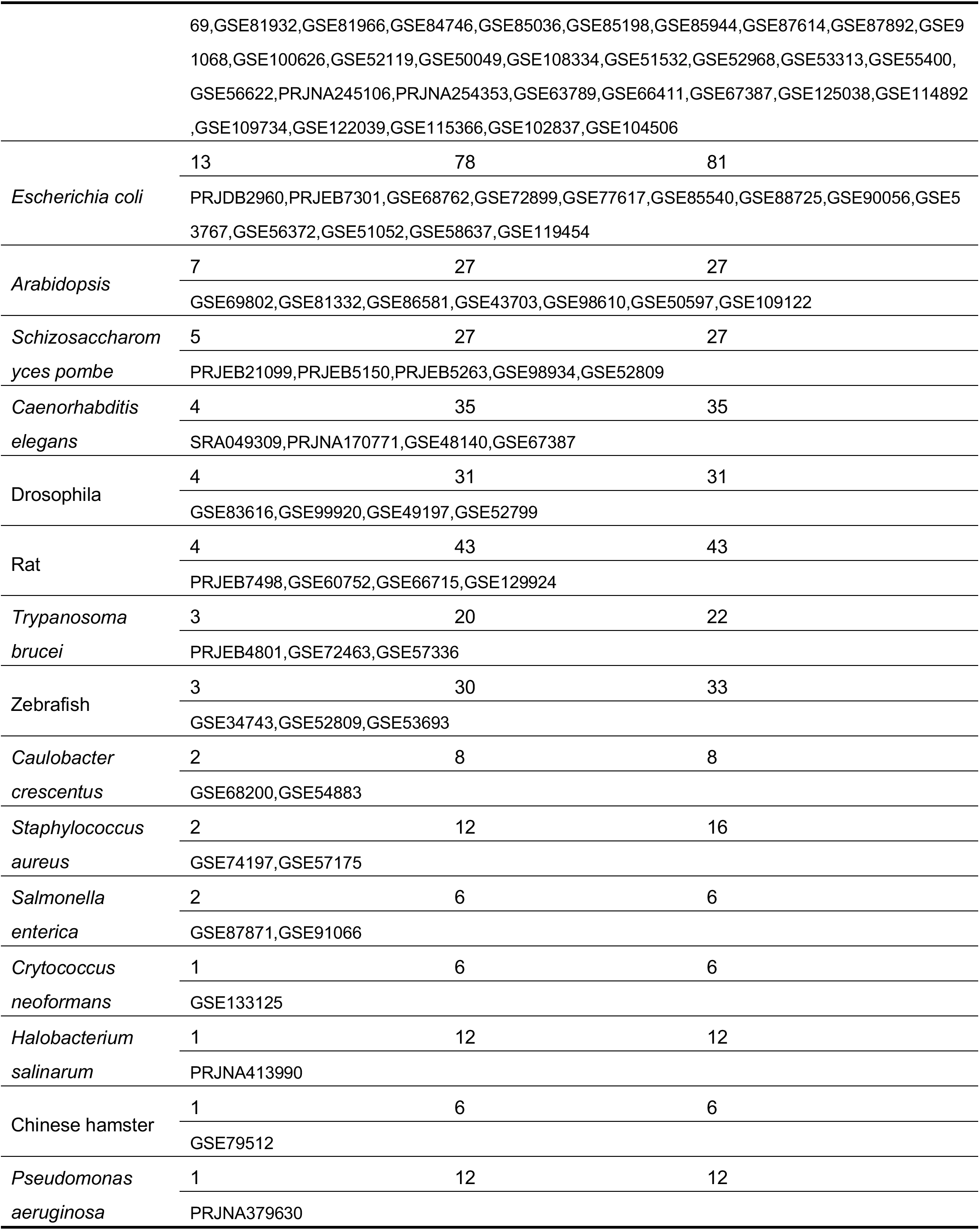
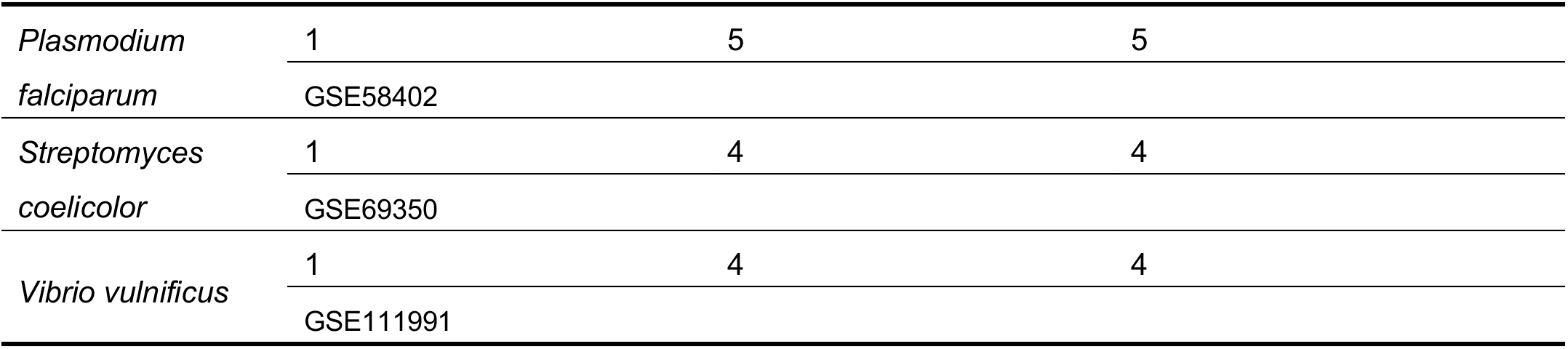
Summary of Ribo-seq and matched RNA-Seq datasets used in this study.

**Additional file 2: Figure S1.**
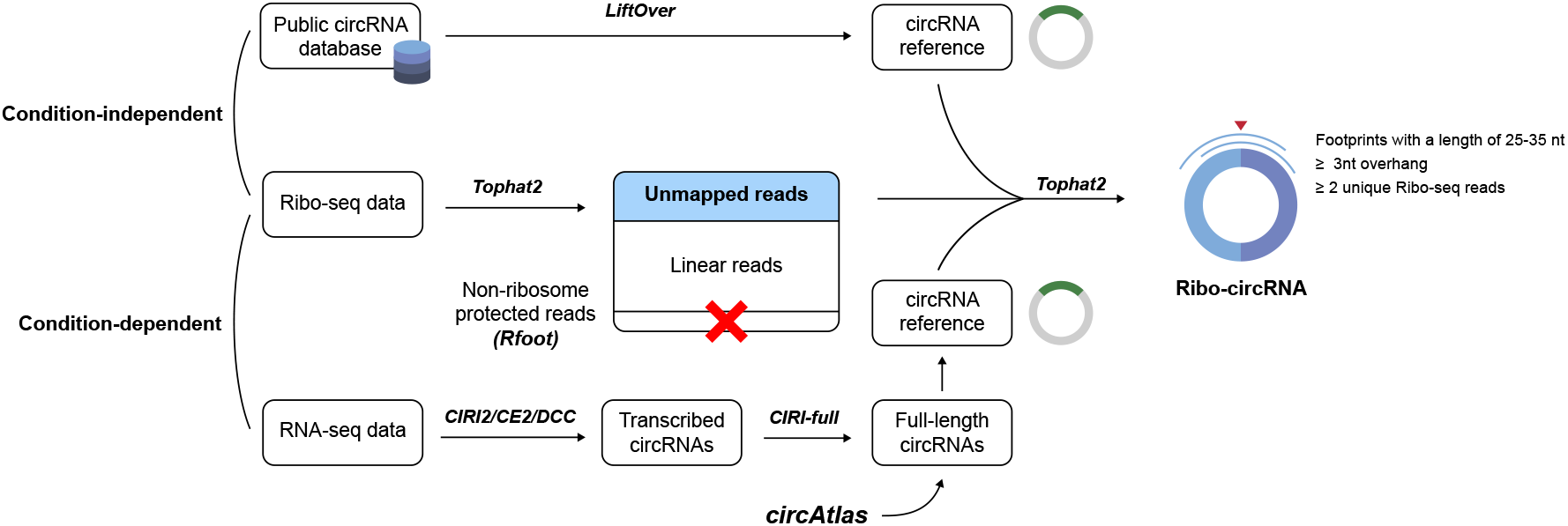
Flow diagram of processing pipeline for translatable circRNAs. Two different strategies were used to characterize ribosome-associated circRNAs: condition-dependent detection for Ribo-seq and perfectly matched RNA-seq datasets and conditionindependent detection for previously reported circRNAs and Ribo-seq datasets.

**Additional file 3: Table S2.** Summary of circRNAs reported in the public databases.

| Database | Species | Genome | LiftOver | Number of circRNAs | Pubmed ID | Note |
| --- | --- | --- | --- | --- | --- | --- |
| circAtlas | Human | hg38 | hg38 | 580,718 | 32345360 | - |
|  | Mouse | mm10 | mm10 | 252,811 |  |  |
|  | Rat | rn6 | rn6 | 97,378 |  |  |
| circBank | Human | hg19 | hg38 | 138,414 | 31023147 | - |
| circBase | Human | hg19 | hg38 | 138,477 | 25234927 | - |
|  | Mouse | mm9 | mm10 | 16,420 |  |  |
|  | C.elegans | ce6 | ce11 | 718 |  |  |
|  | Drosophila | dm3 | dm6 | 3,406 |  |  |
| CIRCpedia | Human | hg38 | hg38 | 183,993 | 30172046 | - |
|  | Mouse | mm10 | mm10 | 55,312 |  |  |
|  | Rat | rn6 | rn6 | 10,197 |  |  |
|  | C.elegans | ce10 | ce11 | 3,840 |  |  |
|  | Drosophila | dm6 | dm6 | 8,560 |  |  |
|  | Zebrafish | danRer10 | danRer11 | 891 |  |  |
| circRNADb | Human | hg19 | hg38 | 32,472 | 27725737 | - |
| CSCD | Human/cancer | hg38 | hg38 | 507,197 | 29036403 | - |
| exoRBase | Human/blood<br>exosomes | hg38 | hg38 | 58,330 | 30053265 | - |
| TSCD | Human | hg38 | hg38 | 128,470 | 27543790 | - |
|  | Mouse | mm10 | mm10 | 5,038 |  |  |
| Circ2Disease | Human | hg19 | hg38 | 248 | 30030469 | - |
| MiOncoCirc | - | - | - | - | 30735636 | incomplete annotation |
| Circad | - | - | - | - | 32219412 | lack of a download link |
| CircFunBase | - | - | - | - | 30715276 | lack of a download link |
| CircR2disease | - | - | - | - | 29741596 | incomplete annotation |
| CircRNADisease | - | - | - | - | 29700306 | incomplete annotation |
| LncRNADisease | - | - | - | - | 23175614 | incomplete annotation |
| CircInteractome | - | - | - | - | 26669964 | lack of a download link |
| CircNet | - | - | - | - | 26450965 | Inaccessible webpage |

**Additional file 4: Table S3.**
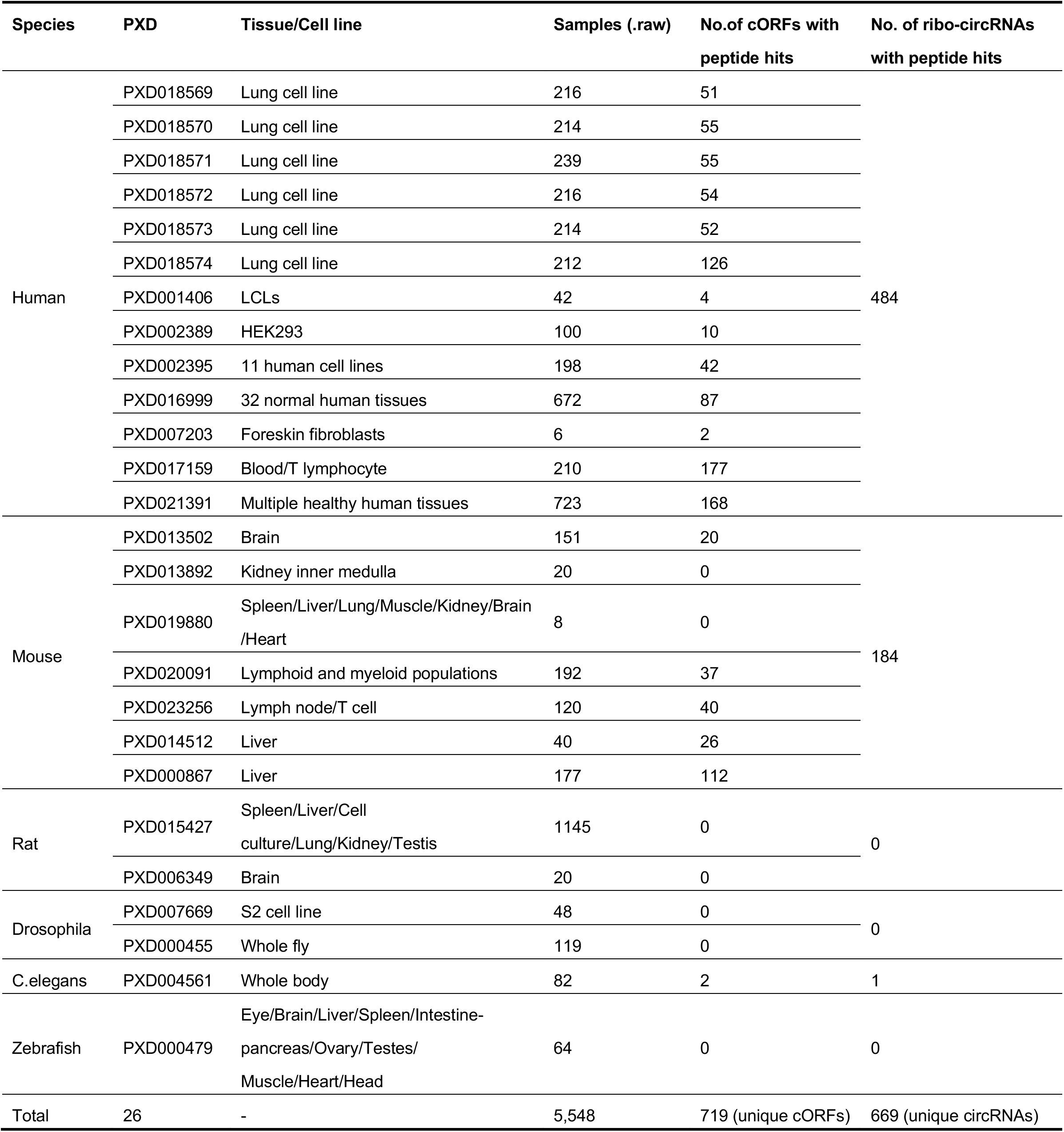
Summary of public proteomics datasets used in this study.

